# Kinetics Of Interferon-λ And Receptor Expression In Response To *In Vitro* Respiratory Viral Infection

**DOI:** 10.1101/2021.12.30.474514

**Authors:** Alexey A Lozhkov, Nikita D Yolshin, Irina L Baranovskaya, Marina A Plotnikova, Mariia V. Sergeeva, Natalia E Gyulikhandanova, Sergey A Klotchenko, Andrey V Vasin

## Abstract

The major protective immune response against viruses is production of type I and III interferons (IFNs). IFNs induce the expression of hundreds of IFN-stimulated genes (ISGs) that block viral replication and further viral spread. The ability of respiratory viruses to suppress induction of IFN-mediated antiviral defenses in infected epithelial cells may be a factor contributing to the particular pathogenicity of several strains. In this report, we analyzed expression of IFNs and some ISGs in an alveolar epithelial cell subtype (A549) in response to infection with: influenza A viruses (A/California/07/09pdm (H1N1), A/Texas/50/12 (H3N2)); influenza B virus (B/Phuket/3073/13); adenovirus type 5 and 6; or respiratory syncytial virus (strain A2). *IFNL* and ISGs expression significantly increased in response to infection with all RNA viruses 24 hpi. Nevertheless, only IBV led to early increase in *IFNL* and ISGs mRNA level. IBV and H1N1 infection led to elevated proinflammatory cytokine production. We speculate that augmented IFN-α, IFN-β, IL-6 levels negatively correlate to *SOCS1* expression. Importantly, we showed a decrease in *IFNLR1* mRNA in case of IBV infection that implies the existence of negative ISGs expression regulation at IFNλR level. It could be either a specific feature of IBV or a consequence of early *IFNL* expression.

## Introduction

In response to viral infection, components of the innate immune response are activated (Cole and Ho, 2017). The most important components of the innate immune response are type I and III interferons (IFNs) (Cole and Ho, 2017). IFNs induce activation of defense mechanisms and prepare cells for possible viral invasion. While the antiviral properties of type I IFNs have been widely studied (Randall and Goodbourn, 2008), much less is known about the features of type III IFNs (IFN-λ). In humans, four IFN-λ subtypes have been found: IFN-λ_1_ (IL-29); IFN-λ_2_ (IL-28A); IFN-λ_3_ (IL-28B); and IFN-λ_4_. IFN-λ are encoded by *IFNL*1- 4 genes. Among IFN-λ_1-3_, there is high conservation of amino acid sequence (Miknis et al., 2010). The actions of IFN-λ on the cell are carried out by binding to the heterodimeric receptor (IFNλR). IFN-λ functions significantly overlap with those of type I IFNs and induce the expression of analogous interferon-stimulated genes (ISGs) (Crotta et al., 2013). Expression of both type I and type III IFNs is induced by the activation of the two most important cytosolic sensors, retinoic acid-inducible gene I (RIG-I) and melanoma differentiation-associated protein 5 (MDA5). RIG-I and MDA5 appear to differentially stimulate IFNs in response to different virus-derived structures, with RIG-I generally responding most potently to 5′ di and tri-phosphate double-stranded RNAs (dsRNA); MDA5 preferentially associates with long dsRNA (Brisse and Ly, 2019).

Despite obvious similarities in mechanisms of induction and downstream signaling, there are obviously some differences in the functioning of type I and type III IFNs. Presumably, type I IFNs have the potential to induce inflammation in addition to antiviral function, while type III IFNs promote the production of antiviral ISGs without the function of inducing inflammation (Sun Y et al., 2018).

The selectivity of type III IFNs is due to peculiarities of receptor subunit expression. IFN-λ actions are carried out through the heterodimeric IFNλR receptor, consisting of the IFNλR1 and IL10R2 subunits. The IL10R2 subunit is also part of the receptor complexes for IL-10, IL-22, and IL-26; it is expressed in cells of various tissues (Miknis et al., 2010). Expression of the IFNλR1 subunit demonstrates a more limited cellular distribution and is present in epithelial cells (Sommereyns et al., 2008), keratinocytes (Zahn et al., 2011) differentiated dendritic cells (Yin et al., 2012; Zhang et al., 2013) and hepatocytes (Dickensheets et al., 2013).

Consequently, the mucous membranes of the respiratory and gastrointestinal tracts are tissues that are mainly targeted by IFN-λ (Sommereyns et al., 2008). This tissue specificity correlates with IFN-λ antiviral activity, which manifests itself mainly in relation to viruses with high tropism for epithelial tissues (Hermant and Michiels, 2014; Lozhkov et al., 2020). This class of viruses includes respiratory viruses such as influenza A and B virus (IAV, IBV), respiratory syncytial virus (RSV), and some types of adenovirus (AdV).

It is well known that respiratory viruses induce IFNs and ISGs production. However, a vast majority of research is focused on features of one or several viral strains, meanwhile matching the data from unrelated research that were carried out in different cell lines should be approached with caution. The number of works that are devoted to direct comparison of the kinetics of IFNs expression stimulated by a wide panel of respiratory viruses is limited. In present research paper we evaluated the dynamics of *IFNL* expression using A549 cells infected with RNA-viruses (IAV, IBV, RSV) and DNA-virus (AdV). AdV and RNA-viruses are quite different in pathogenesis, so that we observed distinct *IFNL* expression.

## Materials and Methods

### Viruses

All viral strains were obtained from the Virus and Cell Culture Collection of the Smorodintsev Research Institute of Influenza (St. Petersburg, Russia). Influenza viruses were grown in 11-day-old embryonated eggs, purified by sucrose gradient, and stored at −80°C. The infectivity values of the viral stocks in MDCK cells were: 3.2×10^7^ TCID_50_/ml for A/California/07/09 ((A)H1N1pdm09); 3.2×10^7^ TCID_50_/ml for A/Texas/50/12 ((A)H3N2); and 3.2×10^5^ TCID_50_/ml for B/Phuket/3073/13 (Yamagata lineage).

Adenovirus working stocks were generated by infecting A549 cells at a multiplicity of infection (MOI) of 0.001 for 72 h. Supernatant was then clarified by centrifugation, aliquoted, and stored at −80°C. The infectivity of the viral stocks in A549 cells was 3.2×10^6^ TCID_50_/ml for both serotype 5 (AdV-5) and serotype 6 (AdV-6). The RSV A2 strain was grown in HEp-2 cells. The infectivity of the RSV stocks in HEp-2 cells was 3.2×10^7^ TCID_50_/ml.

### Infection of Cells

The A549 (CCL-185) cell line was obtained from the American Type Culture Collection (ATCC). A549 cells (human type II alveolar epithelial line) were cultured in F12K medium (Gibco, USA) supplemented with 10% fetal bovine serum (Gibco, USA). For infection, cells were seeded onto 12-well plates (Thermo Scientific Nunc, USA) at 5×10^5^ cells per well. Around 100% confluent monolayers were washed with DPBS (Gibco, USA) and infected at a multiplicity of infection (MOI) of 1. After 60 min of adsorption at 37°C, virus-containing inoculum was removed, and 1 ml of fresh medium was added. Every plate contained at least three replicates of uninfected cells. The infected cells and non-infected controls were incubated at 37°C (5% CO_2_ with humidification) and harvested at 4, 8, and 24 hours after infection.

### Primer and Probe Design

Primers and fluorescent oligonucleotide probes, containing fluorescent reporter dyes at the 5’-end and a quencher at the 3’-end (Table 1), were commercially synthesized and HPLC-purified (Evrogen, Russia).

**Table 1.**
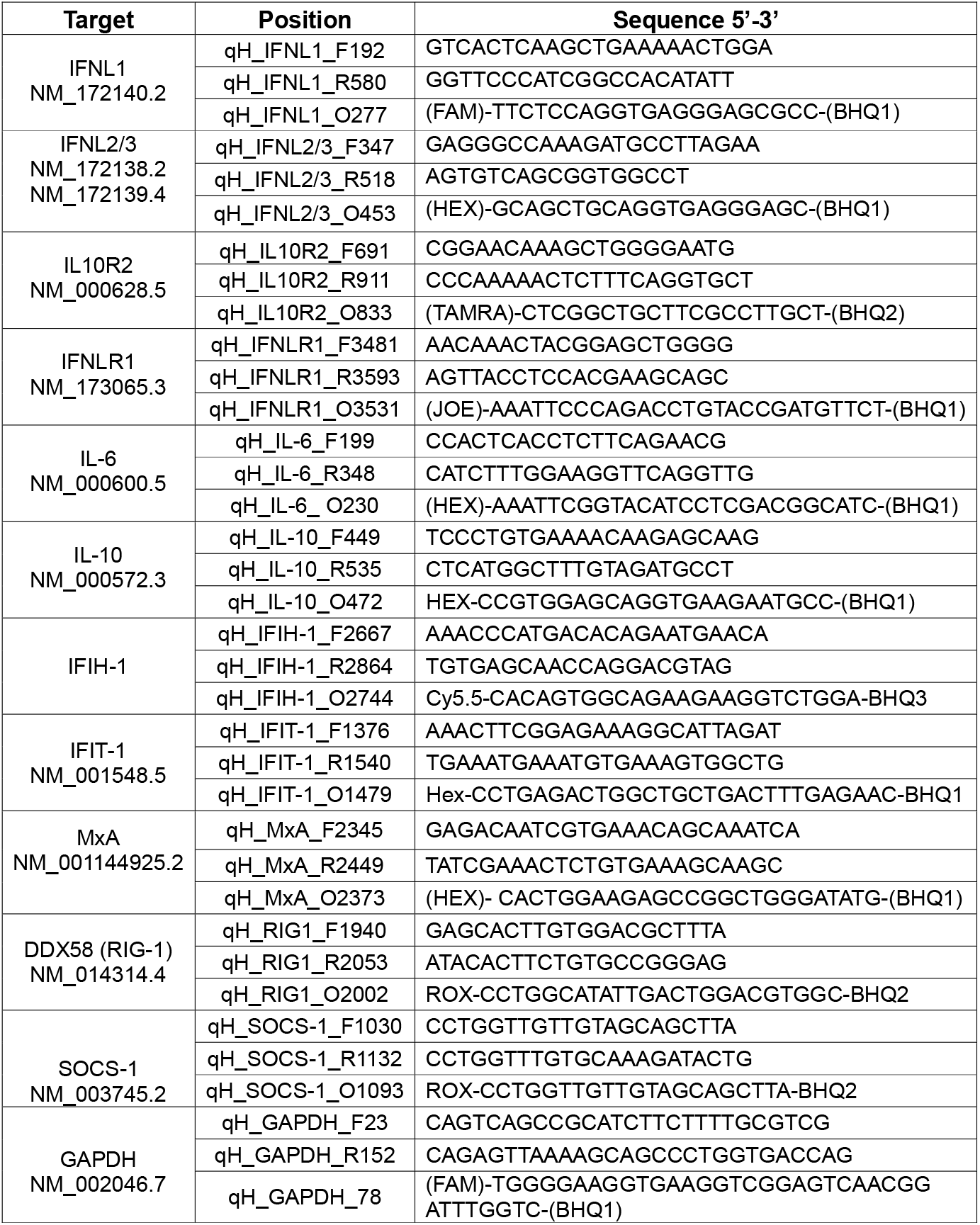
Primers and probes.

### RNA isolation

Total RNA was isolated from A549 cells using TRIZol reagent (Invitrogen, USA) according to the manufacturer’s instructions. RNA concentrations and integrity were analyzed using a NanoDrop ND-1000 spectrophotometer (NanoDrop Technologies, USA).

### Reverse Transcription Reaction

Two micrograms of total RNA were treated by DNase (Promega) and then directly reverse transcribed using oligo-dT_16_ primers and MMLV reverse transcriptase (Promega). Complementary DNA synthesis was carried out at 42°C for 60 min; products were stored at −20°C until use.

### PCR analysis

Real-time PCR assays were performed using the CFX96 Real-Time PCR System (Bio-Rad, USA). Evaluation of *IFNL*1, *IFNL*2-3, *IL10RB, IFNLR1, MxA, IFIT1, RIG-1, MDA5, IFNB and SOCS1* genes expression was performed in 25 μL reaction solutions containing 12.5 μL BioMaster HS-qPCR mix (2x) (BioLabMix, Russia), 2 μL cDNA (diluted 1:2), and 0.4 μM of each primer and probe.

For viral RNA gene expression analyses, the CDC Influenza A/B Typing Kit (# FluIVD03-1, Centers for Disease Control and Prevention, USA) and the Real-time RT-PCR Assay for RSV (Centers for Disease Control and Prevention, USA) were used.

### ELISA

Human IFN-λ_1/3_, IL-6, and IL-10 concentrations in cell culture supernatants were measured by enzyme-linked immunosorbent assay (ELISA) using commercial kits or antibodies. Lambda IFN levels were evaluated by human IL-29/IL-28B (IFN-lambda 1/3) DuoSet ELISA (DY1598B, R&D Systems, USA). IL-6 was measured by human IL-6 DuoSet ELISA (DY206, R&D Systems, USA). Measurement of IL-10 was performed using: Rat Anti-Human IL-10 Capture Antibodies (554705, BD Biosciences, USA); Biotin Anti-Human and Viral IL-10 Detection Antibodies (554499, BD Biosciences, USA); and Recombinant Human IL-10 (1064-IL-010, R&D Systems, USA). Viral loads were evaluated using in-house antibodies (Plotnikova et al., 2020).

### Statistical data processing

Data processing was carried out in Microsoft Excel. GraphPad Prism was used to evaluate the statistical significance of differences.

## Results

### Interferon response of A549 cells to infection with respiratory viruses

In our study, we examined cellular immune responses to infection of A549 epithelial cells with RNA viruses (IBV, IAV H1N1pdm09, IAV H3N2, RSV) and a DNA virus (AdV serotype 5 and 6) at the same MOI. The replicative cycle of influenza is about 8 hours. The production of mRNA and proteins of RSV reaches its peak by 15-20 hpi, while AdV life cycle is a bit longer. Cells were infected without trypsyn to exclude the possibility of infection with viral offspring. So, IFNs and ISGs production refers to a single replicative cycle of any virus. We choose 4 hpi, 8 hpi, and 24 hpi timepoints for comparison of *IFNL* expression kinetics. At first, we showed that the viral genomes are capable of effectively replicating in the A549 cell culture selected (Supplementary Materials, Figure S1). Hence, all of the selected viruses exhibited the ability to replicate and to form new viral particles in A549 cells.

Changes in IFN expression were virus specific. Significant increases in *IFNL* (1, 2/3) as well as IFNB mRNA levels were observed upon infection with all RNA viruses (Figure 1a,b). Increases in *IFNL* expression correlated with accumulation of RNA from these viruses. The kinetics of *IFNL* expression, in response to IBV infection, was fundamentally different compared to the other RNA viruses. Already at 4 hpi, an increase in *IFNL* level, by more than 10,000-fold, was observed compared with the control.

**Figure 1:**
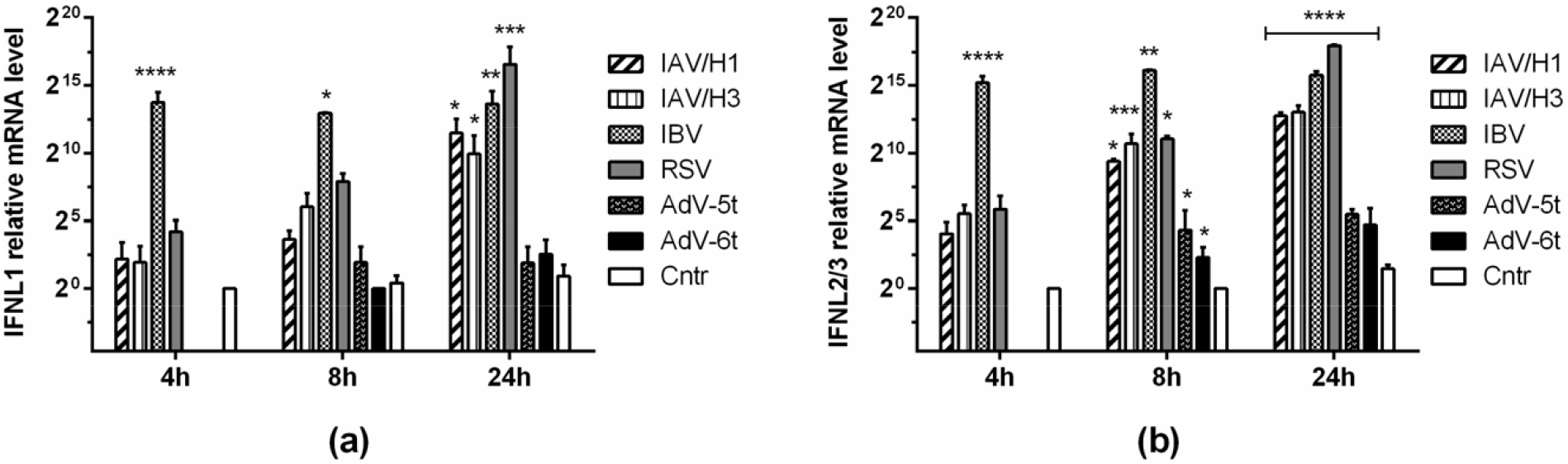
Respiratory viruses stimulate IFN-λ gene expression with different kinetics. Expression levels of IFNL (a) and IFNL2/3 (b) mRNA were measured at 4, 8, and 24 hpi. Gene expression was analyzed via ΔΔCt method (relative to GAPDH). Statistical significance (p-value) was determined by ordinary one-way ANOVA, followed by a pairwise Dunnett’s multiple comparisons test: **** ― Adjusted P Value < 0.0001; *** ― < 0.001; ** ― < 0.01; * ― < 0.05 compared to Cntr. Cntr – intact cells that were cultured in the same conditions and were not infected (instead, sterile medium F12K was added). At least three biological replicates were used for each experimental data point. Data are represented as mean ± SD.

IFN-λ protein level in cell culture supernatants was assessed. It was found that a significant increase in IFN-λ is observed only with IBV or RSV infection (Figure 2a). The levels of IFN-α and IFN-β increased in response to IBV and (A)H1N1pdm09 (Figure 2b,c). Despite the fact that IFNs are the primary, universal link in the activation of the innate immune response, we have shown that stimulation of the type I and type III IFN systems is not only virus specific, but also strain specific.

**Figure 2:**
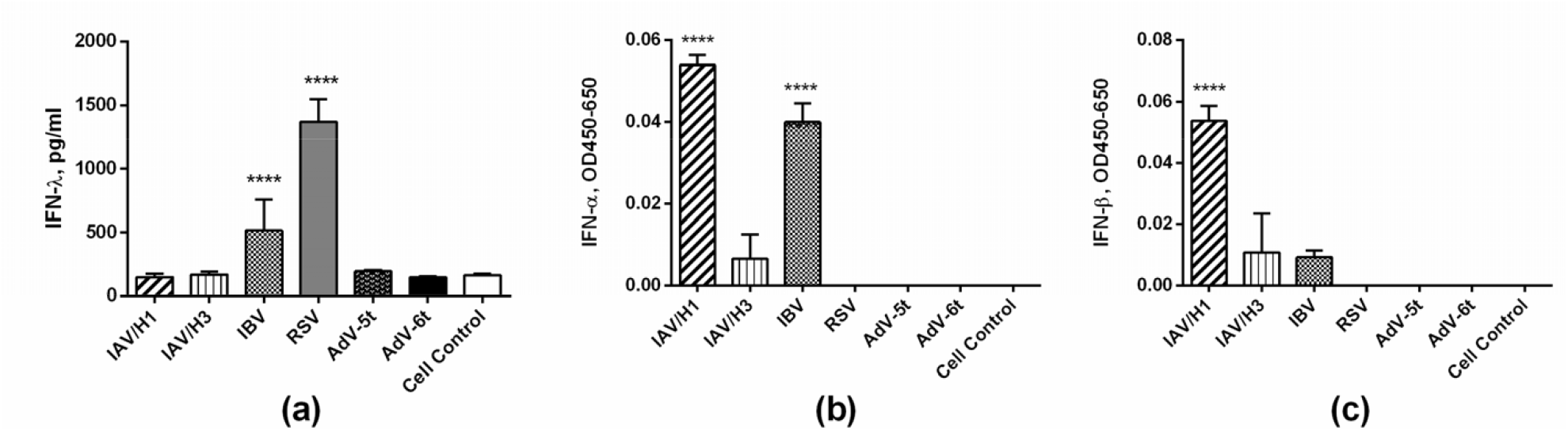
Production of type I and type III IFNs in response to viral infection. Levels of IFN-λ (a), IFN-α (b), and IFN-β (c) were measured by ELISA of cell supernatants 24 hours post infection Statistical significance (p-value) was determined by ordinary one-way ANOVA, followed by a pairwise Dunnett’s multiple comparisons test: **** ― Adjusted P Value < 0.0001 compared to Cntr. Cntr – intact cells that were cultured in the same conditions and were not infected (instead, sterile medium F12K was added). At least three biological replicates were used for each experimental data point. Data are represented as mean ± SD.

### ISG expression

We also evaluated changes in the expression of several ISGs in response to viral infection. In both virus-infected and uninfected cells, IFNs induce the expression of myxovirus resistance protein (MxA), which makes MxA an excellent marker for detecting activation of an IFN-dependent response (Haller et al., 2015). We evaluated *MxA* expression kinetics at 4, 8, and 24 hpi (Figure 3a). In general, *MxA* expression by 24 hpi increased significantly upon infection with all viruses. However, the *MxA* mRNA level in AdV infected cells was significantly lower than those with RNA virus infections. At 4 hpi with IBV, a significant increase in *MxA* expression (about two orders of magnitude) was noted. By 8 hpi with IBV, *MxA* expression had reached its maximum and was also significantly increased compared to all other groups. By 24 hpi with RSV, there was a significant increase in *MxA* expression.

**Figure 3:**
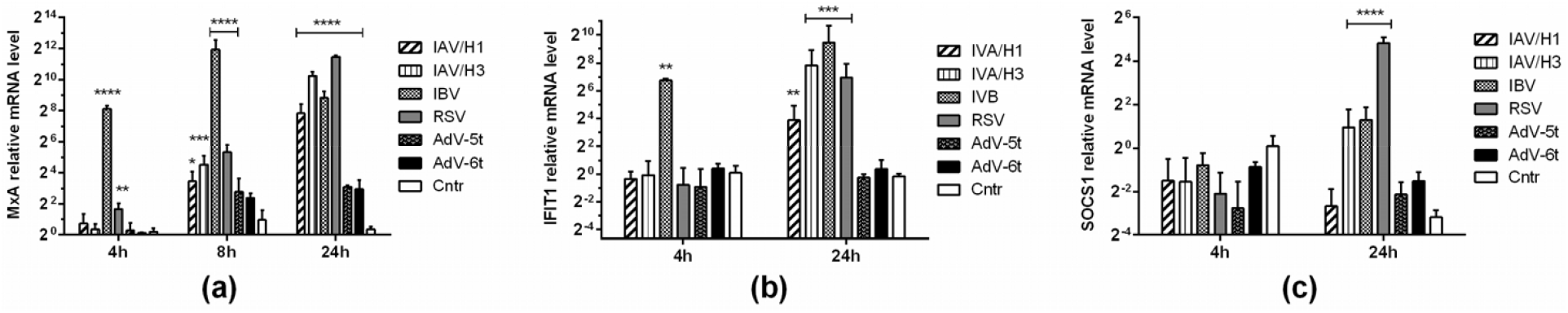
Respiratory viral infection of A549 cells leads to an increase in the expression of classic ISGs: MxA (a); IFIT1 (b); and SOCS1 (c). Gene expression was analyzed via ΔΔCt method (relative to GAPDH). Statistical significance (p-value) was determined by ordinary one-way ANOVA, followed by a pairwise Dunnett’s multiple comparisons test: **** ― Adjusted P Value < 0.0001; *** ― < 0.001; ** ― < 0.01 compared to Cntr. Cntr – intact cells that were cultured in the same conditions and were not infected (instead, sterile medium F12K was added). At least three biological replicates were used for each experimental data point. Data are represented as mean ± SD.

The expression pattern for *IFIT1* looked similar (Figure 3b). However, with AdV infection, no significant change in *IFIT1* expression was observed. Therefore, it can be concluded that the increase in *IFNL* expression was largely synchronized with increased *MxA* and *IFIT1* mRNA levels.

We also measured the expression of *SOCS*1 (Figure 3c), which is an inducible negative regulator of IFN lambda. By 4 hpi, we did not find any significant changes in expression. By 24 hpi, however, *SOCS*1 expression was significantly increased in cells when infected with (A)H3N2, IBV, or RSV.

### Assessment of proinflammatory cytokine and chemokine levels

The levels of specific cytokines (IL-6, IL-8, IL-10) in cell culture supernatants were analyzed (Figure 4). IL-6 levels were significantly increased with IBV and (A)H1N1pdm09. The IL-6 levels for RSV and (A)H3N2 also exceeded control values. Differences in IL-10 levels were found for IBV and (A)H1N1pdm09. With the (A)H3N2, RSV, and AdV5 viruses, infection led to an increase in the level of chemokine IL-8. Thus, secreted levels of IL-6 and IL-10 generally correlated with type I IFN production. A significantly increased IL-8 level was a specific feature of (A)H3N2 infection in A549 cells.

**Figure 4:**
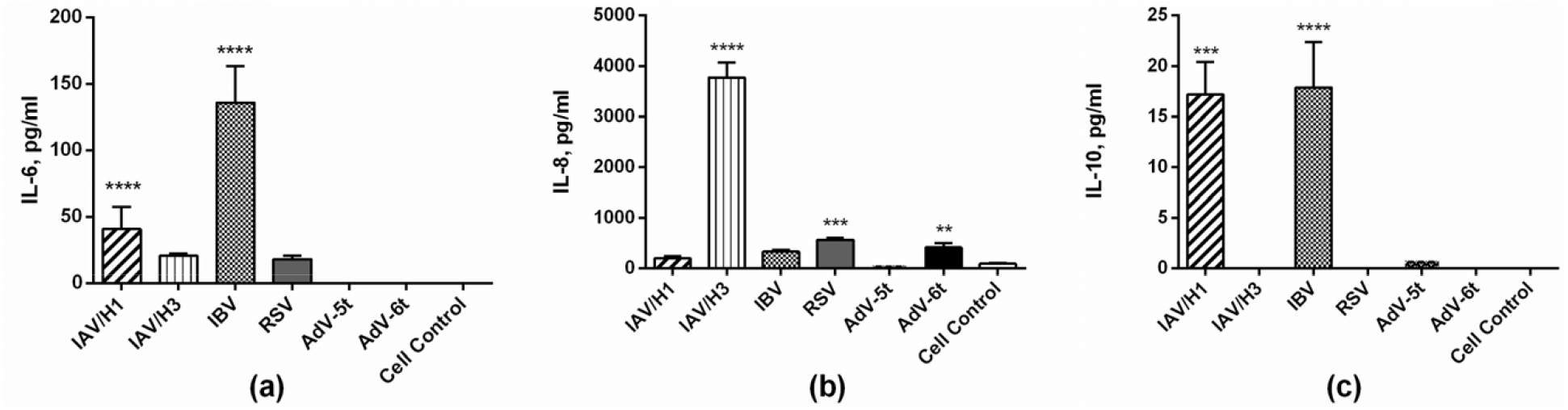
A549 cell cytokine production patterns differ when infected with various viral strains. IL-6 (a), IL-8 (b), and IL-10 levels were measured by ELISA in cell supernatants 24 hours post infection. Statistical significance (p-value) was determined by ordinary one-way ANOVA, followed by a pairwise Dunnett’s multiple comparisons test: **** ― Adjusted P Value < 0.0001; *** ― < 0.001; ** ― < 0.01 compared to Cntr. Cntr – intact cells that were cultured in the same conditions and were not infected (instead, sterile medium F12K was added). At least three biological replicates were used for each experimental data point. Data are represented as mean ± SD.

### Determination of expression of IFNλR subunits and the cytosolic sensors MDA5 and RIG- 1

It should be noted that *MDA5* and *RIG-1* expression is induced by autocrine or paracrine action of IFNs, so that an increase in mRNA level of these genes can be considered as positive feedback loop that could further augments IFNs production. Already by 4 hpi, IBV infection led to an increase in the expression of both the *RIG*-1 and *MDA5* cytosolic sensors (Fig. 5). By 24 hpi, *RIG*-1 expression was also significantly increased in response to infection with IBV, A/H3N2, or RSV. Notably, an increase in *MDA5* expression was observed for all RNA viruses.

**Figure 5:**
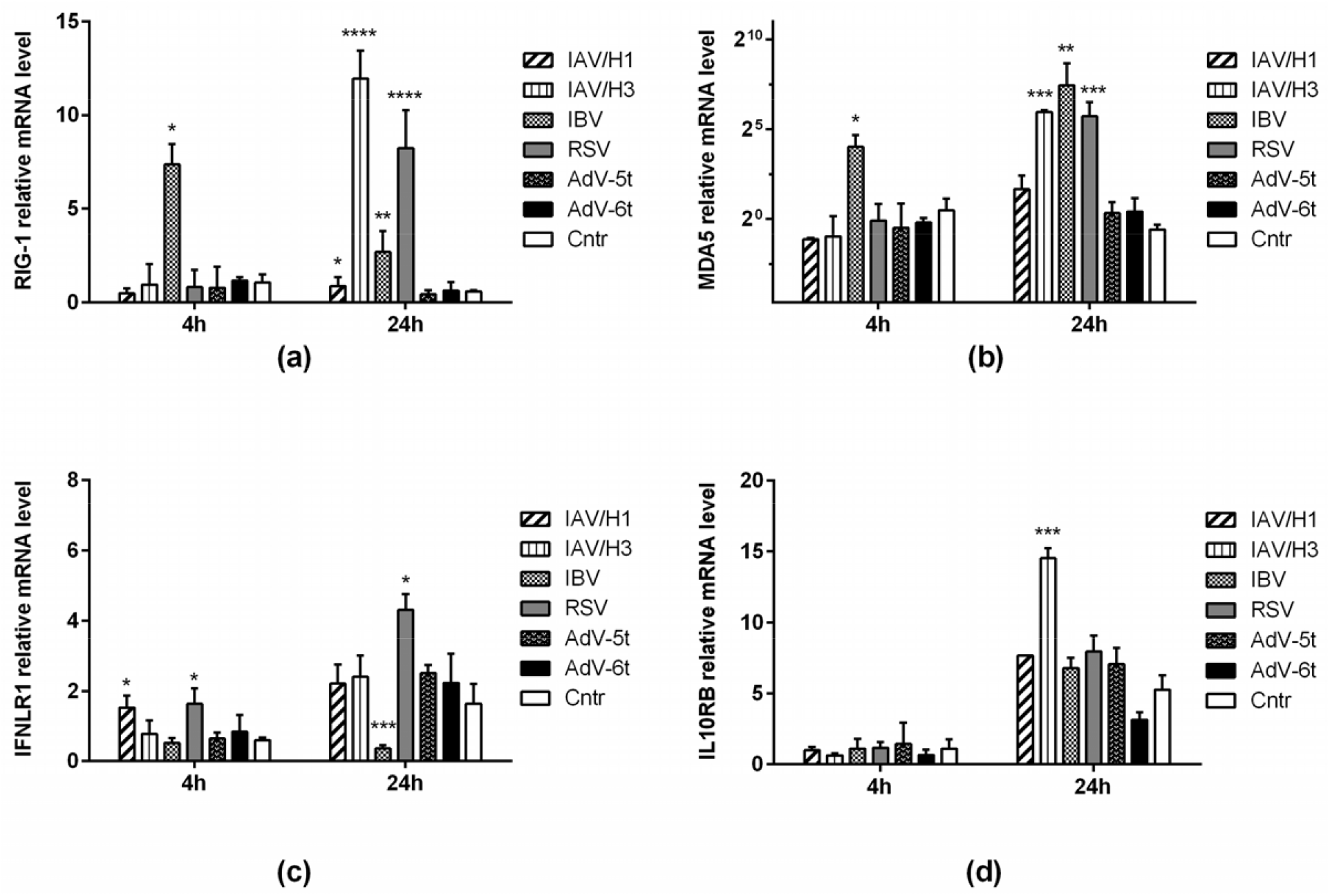
The expression kinetics of IFN response signal transduction genes differ in A549 cells infected with various respiratory viruses. Gene expression was analyzed via ΔΔCt method (relative to GAPDH). The expression of IFNL receptor subunits IFNLR1 (a) and IL10RB (b), as well as the cytosolic RLRs Rig-1 (c) and MDA5 (d), were assessed by RT-PCR. Statistical significance (p-value) was determined by ordinary one-way ANOVA, followed by a pairwise Dunnett’s multiple comparisons test: **** ― Adjusted P Value < 0.0001; *** ― < 0.001; ** ― < 0.01 compared to Cntr. Cntr – intact cells that were cultured in the same conditions and were not infected (instead, sterile medium F12K was added). At least three biological replicates were used for each experimental data point. Data are represented as mean ± SD.

Possible regulation of IFN-λ-dependent signaling activation by variation in IFNλR subunit expression was evaluated. The expression levels of *IFNLR1* and *IL10R2* were assessed. A significant decrease in the expression of *IFNLR1* subunit (more than 5-fold compared to the control) was noted one day after IBV infection (Figure 4). Presumably, the decrease in *IFNLR1* level was associated with a rapid increase in *IFNL* expression. There were no significant changes in the expression levels of *IL10R2*, a nonspecific IFNλR subunit.

## Discussion

In our previous work (Plotnikova et al., 2021) we showed that IFN-λ_1_ exhibits antiviral activity against various RNA viruses (IAV, SARS-CoV-2, CHIKV). IFNs are a major component of innate defense against viruses. The production of endogenous IFN-λ by epithelial cells is a natural defense mechanism that limits the growth and spread of RNA viruses. In this work, we assessed the dynamics of *IFNL* and several ISGs expression in response to infection of A549 cells with respiratory viruses (H1pdm09, H3, IBV, RSV, AdV types 5 and 6). In present study it was shown that stimulated by IBV early induction of *IFNL* and ISGs expression is associated with a decrease in mRNA level of *IFNLR1*, the specific subunit of IFNλR.

It is known that the induction of IFNs, proinflammatory cytokines, and chemokines is associated with strain pathogenicity (Cole and Ho, 2017). When studying the IFN status of A549 cells, we showed that infection with RNA viruses led to a significant increase in mRNA, meanwhile AdV infection elicited only a weak increase in *IFNL* and *IFNB* mRNA (Figure 1). With IBV, type III IFNs were extremely elevated, with a peak at 4 hpi. It has been observed, in monocyte-derived dendritic cells, that IBV induces early expression of *IFNL*1 and *IFNB* mRNA (as early as 2 hpi). IAV causes noticeable activation of these genes much later (only starting from 8-12 hpi) (Strengell et al., 2012; Sun Y et al., 2018). Differences in IFN expression kinetics, obtained here for IAV and IBV, on the whole agree with results described in the literature. In turns, AdV can evade the early IFN-dependent immune response.

In our work, we used AdVs (types 5 and 6) which belong to serotype C and exhibit high tropism for respiratory epithelial cells (Chahal et al., 2012). Despite the absence of significant changes in *IFNL*1 expression, by 24 hours for both AdV5 and AdV6, an increase in *IFNL*2/3 mRNA (4 to 6-fold) was observed (Figure 1). In the modern literature, it has been shown that AdV has a suppression system for the IFN-induced antiviral response (Chahal et al., 2012).

We performed a qPCR analysis to determine the expression levels (4 and 24 hpi) of antiviral ISGs: *MxA* and *IFIT1*. Already by 4 hpi, both *MxA* and *IFIT1* displayed markedly elevated mRNA levels with IBV infection. The level of *IFIT1* mRNA significantly increased only when cells were infected with RNA viruses (Figure 3). In general, these ISGs profile was synchronized with *IFNL* expression. This way, we observed distinct *IFNL* and ISGs expression in case of the different respiratory viruses. Next, we aimed to define the regulatory factors that could influence on *IFNL* and ISGs expression profile.

It has been shown that IAV infection induces *IFNL* expression mainly through RIG-I-dependent pathway (Wei et al., 2014). Induction of IFN expression occurs already in response to IBV penetration into the cell; and RIG-I cytosolic RNA sensors play a key role in virus recognition (Mäkelä et al., 2015). Presuming that both viral genome replication and the production of IFNs can lead to a change in the expression of cytosolic sensors by positive feedback mechanisms (*RIG*-I and *MDA5* are also ISGs), we evaluated the expression of both RLRs at an early stage (4 hpi) and a late stage (24 hpi) of infection. Infection with RNA viruses resulted in an increase in *MDA5* and *RIG*-1 expression by 24 hpi, with the exception of IAV H1N1 (Figure 5).

Type I and III IFNs can up-regulate SOCS proteins, which negatively regulate IFN signaling by inhibiting the JAK-STAT signaling pathway (Schneider et al., 2014). Here, we found that *SOCS*1 expression was elevated in IAV H3N2, IBV, or RSV-infected cells 24 hpi. In general, these observations are consistent with the changes in RLR expression (Figure 5). Importantly, *SOCS*1 mRNA level was not increased in case of IBV 4 hpi, whereas expression of other ISGs (*MxA, IFIT1, RIG*-I, *MDA5)* and *IFNL* was clearly elevated at this timepoint. It has been shown that the physiological role of SOCS1 proteins is to prevent tissue damage caused by the potent pro-inflammatory effects of type I IFNs (Blumer et al., 2017). Along these lines, attenuation of *SOCS*1 expression can serve as a marker indicating an increased potential of IAV H1N1 and IBV to cause hypercytokinemia.

According to Sun’s assumptions (Sun Y et al., 2018), type I IFNs have the potential to induce inflammation in addition to antiviral function, while lambda IFNs promote the production of antiviral ISGs without the excessive inflammation. In our study, IBV infection was associated with a cytopathic effect and led to increases in the proinflammatory factors IFN-α, IFN-β, IFN-λ, IL-6, and IL-10. It should be noted that although IL-10 itself cannot be attributed to mediators that promote inflammation, it can be a marker of uncontrolled immunopathology (Guo and Thomas, 2017). With influenza (A)H1N1pdm09, the production kinetics of these cytokines were generally the same, with the exception of IFN-λ. This may be evidence of a cytopathic effect of (A)H1N1pdm09 and IBV that is related to decrease in *SOCS*1 expression.

Modulation of IFN signaling can be accomplished by alteration of receptor subunit expression (Stanifer et al., 2019). For instance, published work has established that in nasopharyngeal swabs of children with a severe course of rhinovirus, *IFNLR1* expression was increased compared to samples of children infected with RSV (Pierangeli et al., 2018). Evaluation of IFNλR subunit expression showed that only with IBV was there a slight decrease in *IFNLR1* mRNA level, while non-specific subunit *IL10R2* mRNA level did not change (Figure 5). At the moment, there is not much information available regarding molecular mechanisms in negative regulation of *IFNLR1* expression (Stanifer et al., 2019). In any case, decreased *IFNLR1* expression appears to be a natural compensatory mechanism realized in response to excessive activation of IFNλR-mediated signaling.

## Conclusion

In present study we compared the expression profile of *IFNL* and several ISGs in response to infection A549 with a panel of widely disturbed respiratory viruses. A549 cells are standard cell line that a huge amount of virological experiments are carried out. Although the study could be considered as summarizing and generalization of previously known data, our work highlighted unique features of type III IFN production that should be taken into account by studies examining viral pathogenicity. We speculate that production of proinflammatory cytokines negatively correlate to *SOCS1* expression. Importantly, we showed a decrease in *IFNLR1* mRNA in case of IBV infection that implies the existence of negative ISGs expression regulation at IFNλR level. It could be either a specific feature of IBV or a consequence of early *IFNL* expression.

## Abbreviations

AdV: adenovirus
IAV: influenza A virus
IBV: influenza B virus
IFN(s): interferon(s)
IFNλR: interferon-λ receptor
ISG(s): interferon stimulated gene(s)
RLR: RIG-I-like receptor(s)
RSV: Respiratory syncytial virus

## Acknowledgments

The authors would like to acknowledge the kind help of Edward S. Ramsay for his assistance (translation, editing) in preparation of this paper.

## Conflicts of Interest

The authors declare that there are no conflicts of interests regarding the publication of this paper.

## Funding

This work was supported by a Russian State Assignment for Fundamental Research (0784- 2020-0023).

## Legends to figures

**Figure S1:**
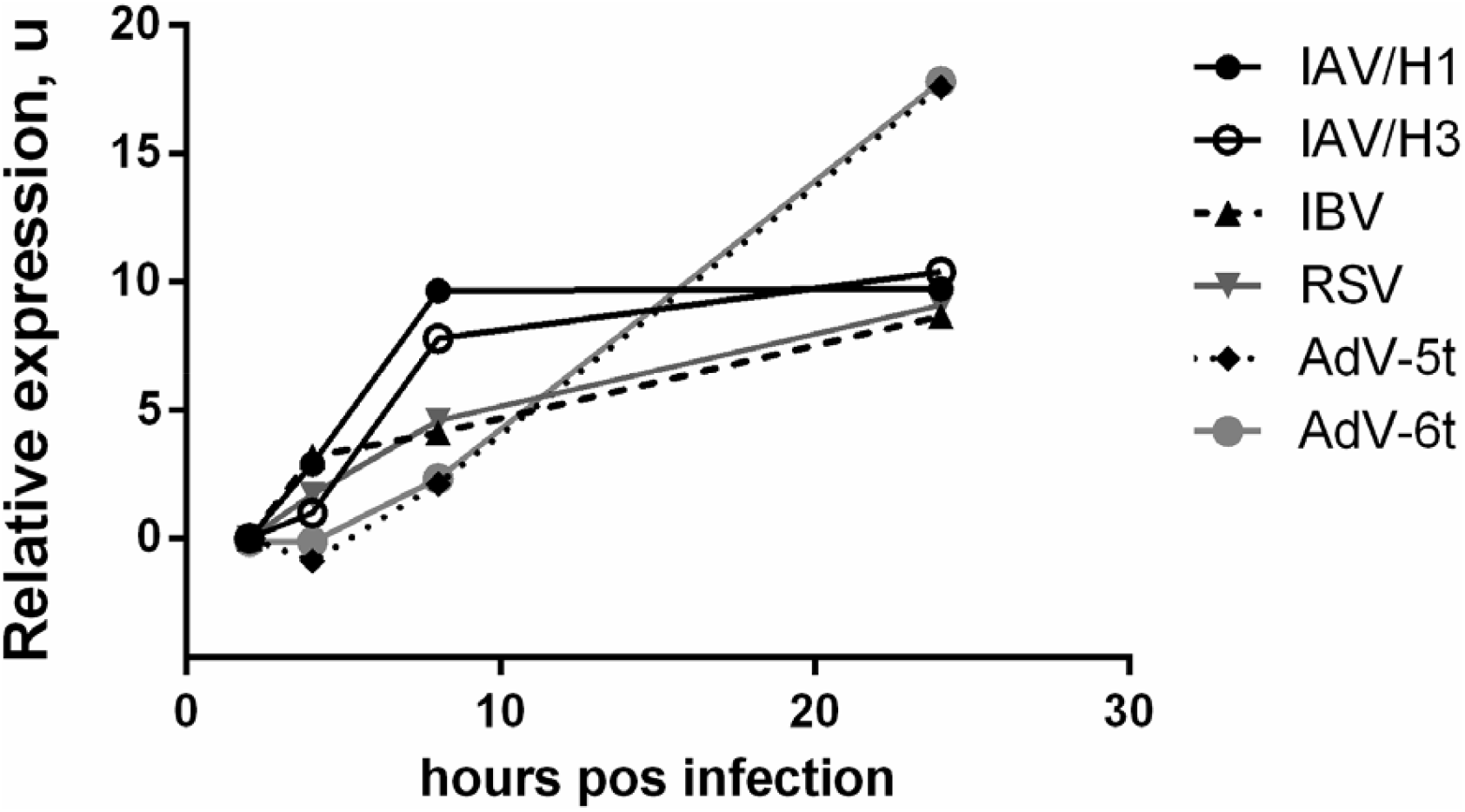
The specified respiratory viruses are capable of replication in A549 cells. Comparison of relative genomic replication rates. The y-axis is the decimal logarithm of viral gene expression. Gene expression was calculated using the ΔΔCt method (relative to GAPDH). When calculating expression, the viral genome expression level at 2 hpi was used for normalization of viral replication at later time points. For example, IAV expression at 4 hpi or later was normalized to IAV expression at 2 hpi, etc. The x-axis shows the time after infection. Cntr – intact cells that were cultured in the same conditions and were not infected (instead, sterile medium F12K was added). At least three biological replicates were used for each experimental data point.

## Notes

### Competing Interest Statement

The authors have declared no competing interest.

